# Charge Based Boundary Element Method with Residual Driven Adaptive Mesh Refinement for High Resolution Electrical Stimulation Modeling

**DOI:** 10.64898/2026.03.11.711201

**Authors:** Derek A. Drumm, Gregory M. Noetscher, Hannes Oppermann, Jens Haueisen, Zhi-De Deng, Sergey N. Makaroff

## Abstract

Accurate transcranial electrical stimulation (TES), electroconvulsive therapy (ECT), and electroencephalography (EEG) forward modeling requires resolving numerical singularities in the charge density near electrodes and tissue interfaces. We present an adaptive mesh refinement (AMR) strategy for the charge based boundary element method (BEM) accelerated by the fast multiple method (BEM-FMM) including electrode and interface singularities. We derive a new error estimator which considers both local and nonlocal contributions of the single-layer potential operator and construct a refinement criterion based on the difference in charge solution across AMR iterations. We evaluate this approach on a 5-layer sphere model and on multiple subject-specific head models derived from the 7-tissue SimNIBS (headreco) and 40-tissue Sim4Life (head40) segmentations, using both voltage-controlled and current-controlled electrode formulations. Through convergence analysis on the white matter and deep hippocampal targets, we find electric fields with relative residual errors below 0.1% and 1% for SimNIBS and Sim4Life models, respectively. Our results indicate that the residual based AMR applied to BEM-FMM leads to numerically stable TES and EEG forward solutions in realistic head models.

## 1 Introduction

Modeling transcranial electrical stimulation (TES) [1], electroconvulsive therapy (ECT) [2], and electroencephalography (EEG) [3] requires accurate numerical solutions of the quasi-static Maxwell equations. Boundary element methods (BEM) are a standard approach for these applications since they represent tissue interfaces exactly without the need for volumetric meshing. Modern BEM methodologies can reliably predict electric fields both at the cortical surface and within deep brain structures [4]. Furthermore, the use of the fast multipole method allows for the acceleration of BEM (the so-called charge based BEM-FMM), enabling the efficient computation of BEM solutions over extremely high-resolution head models [5–8]. Even with fast multipole acceleration, the accuracy of BEM solutions relies on the quality and resolution of the surface meshes. Anatomical head models contain thin layers and conductivity jumps which present major difficulties for modeling cortical stimulation with deep targets; these difficulties are even more pronounced when considering a highly realistic head model with non-manifold tissue layers (the solutions of which are only feasible via charge based BEM-FMM [6]). Moreover, modeling TES electrodes presents an additional numerical challenge in that singularities in the charge solution are generated on the electrode-skin interface. These challenges make adaptive mesh refinement (AMR) a necessary tool for achieving accurate BEM solutions.

Classical AMR strategies were developed for determining numerical solutions for arbitrary PDEs applicable to both finite element method [9, 10] and BEM [11–13]. These often involve the computation of an error estimator, which serves as a criterion to determine which mesh elements to refine. In the work of [14], it was suggested to use the total charge as a criterion for refinement; this resulted in decent convergence in the charge density and mean electric field when considering the entire head model, and has been subsequently applied to various BEM applications [4, 6, 8]. However, we found that using this criterion resulted in unsatisfying convergence when considering the solution localized to the white matter and deep cortical structures (i.e. the hippocampi). Therefore, we sought an alternative approach.

Through mathematical analysis, one can derive analytic residual-based error estimators with guaranteed rates of refinement and convergence [12, 13]. However, applying these estimators as criteria for mesh refinement is nontrivial since they are defined using the Dirichlet boundary data, which is a priori unknown for the charge based formulation of BEM. Instead, we derive a refinement criterion based on the numerical charge solution while leveraging the mathematical theory of residual based estimators. Particularly, we establish energy norm bounds relating the BEM Galerkin residual to the charge residual, and derive an error estimator which incorporates both local and nonlocal contributions of the single-layer potential operator. Utilizing this estimator as a refinement criterion, we model TES and EEG electrodes on a simple 5-layer sphere model and on subject head models with two segmentations: the 7-tissue SimNIBS headreco segmentation [15], and the 40-tissue Sim4Life head40 segmentation [16]. Across these models, we examine the convergence of quantities of interest relevant to TES and EEG modeling. We are particularly interested in the convergence of the mean electric field on deep tissue structures (such as the hippocampi), but also analyze the convergence of electrode currents and total cortical electric fields.

## 2 Materials and methods

### 2.1 Adaptive Charge-Based Boundary Element Fast Multipole Method

In this section, we will briefly review the charge-based boundary element fast multipole method (BEM-FMM) and work towards developing a criterion for mesh refinement through iterative adaptive passes.

#### Charge-based BEM-FMM

Consider a volume with boundary *S* comprising the disjoint cortical surface tissues of interest. The charge-based formulation of BEM is the integral equation for determining the surface charge density *ρ*(**x**) due to the impressed electric field **E***i* (**x**) [8, 17, 18]

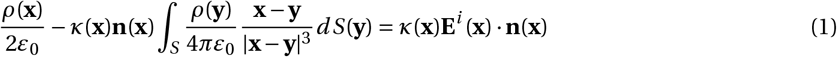

where

- **n**(**x**) is the outward unit normal vector at **x** ∈ *S*;
- *ε*_0_ is dielectric permittivity of free space;
- *κ*(**x**) = (*σ*_−_ −*σ*_+_)/(*σ*_−_ +*σ*_+_) is the conductivity contrast with respect to inner and outer conductivities *σ*_−_ and *σ*_+_, respectively.

Of interest to us in this work is the impressed field induced by a pair of TES or EEG electrodes on the scalp surface, the calculation of which is described in [18]. For a boundary *S* discretized into a triangular mesh, the Galerkin approximation of the charge solution *ρ*_*h*_ of Equation 1 at triangle *T* can be explicitly calculated as [17, 18]

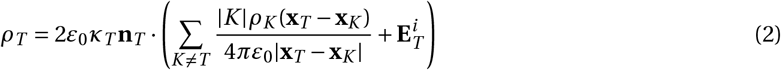

where |*K* | denotes the area of triangle *K*, **x**_*T*_ and **x**_*K*_ the center points of triangles *T* and *K*, respectively, and the calculation of the impressed electric field is accelerated with FMM. Note that here we use shorthand notation to represent values which are implicitly functions of the triangle index (e.g. *ρ*_*T*_ := *ρ*_*h*_ (**x**_*T*_), *κ*_*T*_ = *κ*(**x**_*T*_), etc.). We will use this notation for the remainder of this work whenever clear.

#### Adaptive mesh refinement

We seek a proper choice of refinement strategy as a local estimator of solution error *η* such that if *η*_*T*_ is large enough, then triangle *T* is selected to be refined. Here, we briefly summarize our choice of refinement criterion; full details and derivation may be found in Supplement S1.

We assume that the electric scalar potential *u*(**x**) induced by the surface charge density is a single-layer potential. This single-layer potential can be described as an operator ℒ acting on the charge density. In the discretized context, ℒ is a *N* × *N* matrix whose indices correspond to the boundary triangles *T* in the mesh. Adaptive BEM schemes and convergence rates exist and are well known for the single-layer potential [12, 13], which rely on the computation of the Galerkin residual 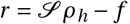 where *f* is Dirichlet boundary data for the potential. In particular, the refinement scheme must ensure that the Galerkin residual *r* and the charge residual *e* = *ρ* − *ρ*_*h*_ vanish in the limit of adaptive passes *l*; that is, ∥*r ∥*_*V*_ − 0 and ∥*e*∥*V* ^∗^ − 0 for some proper choice of function space *V* with corresponding dual *V*^∗^ (see Supplement S1.1). For our problem, the boundary data *f* is a priori unknown (the Dirichlet problem for the potential based BEM has jump conditions on the boundary, see [18]), and thus the Galerkin residual cannot be precisely computed (although it can be approximated, see Supplement S2). Instead, we may bound the Galerkin residual by the charge residual in the energy norm (where *e* is treated as a *N* × 1 vector),

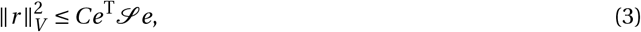

where *C* a positive constant. The discrete operator ℒ possesses scaling properties (see Supplement S1.3) which allow us to split Equation 3 into local and non-local components,

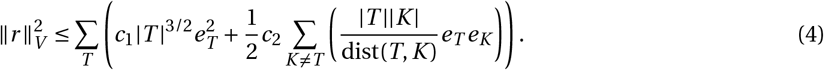

where *e*_*T*_ and *e*_*K*_ denote the evaluation of *e* at triangles *T* and *K*, respectively, and *c*_1_ and *c*_2_ are positive constants.

The right-hand-side of Equation 4 may now act as a criterion for mesh refinement. However, since the true charge *ρ* is unknown, the charge residual *e* is not computable. Therefore, we introduce a surrogate function as a substitute. Denote by *ρ*_*l*_ the Galerkin approximation of the charge and *A*_*l*_ the areas of the mesh triangles (as a function of *T*) at the *l* -th adaptive pass. Then we define the surrogate *ξ*_*l*_ as

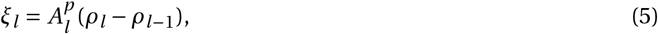

where that *ρ*_*l* −1_ is taken to be the interpolation of the (*l* − 1)-th solution onto the more refined mesh. In Supplement S1.4, we show that this choice of surrogate is sufficient for any *p* ≥ 0 in the sense that the energy norm of *ξ*_*l*_ − *e*_*l*_ vanishes in the limit of adaptive passes. (In practice, we have found that *p* = 3/2 works best as an area scaling, although this may be model dependent). Now, with *ξ*_*l*,*T*_ := *ξ*_*l*_ (**x**_*T*_) denoting the surrogate as a function implicitly dependent on the triangle indices, we define the refinement criterion on the *l* -th adaptive pass as

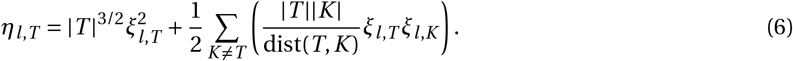

With *η*_*l*,*T*_ serving as a refinement criterion, on each adaptive pass we may use any desired scheme to refine the mesh. For each of the simulations in this work, at adaptive pass *l* we select 5% of the total triangles with the largest *η*_*l*,*T*_ as the target set. These triangles are then refined using barycentric refinement, where each flagged triangle is split into four child triangles each sharing a vertex at the barycenter of the parent triangle.

#### Computation of the refinement criterion

In the initial adaptive pass *l* = 1, clearly our refinement criterion is not computable. There are however a variety of means we can handle this. For instance, we can approximate the “0-th” charge solution *ρ*_0_ as the primary component of the impressed electric field (scaled by the contrast, see Equation 5.6 of [18]), then simply use *ρ*_1_ − *ρ*_0_ as a surrogate on the first step. Another approach is to simply apply a uniform refinement across the entire model on the first adaptive pass, then proceed with *η*_*l*,*T*_ as the refinement criterion. For the simulations shown in this work, this is the approach we take.

Also, for large models, the non-local term in Equation 6 is not so simple to calculate. In the derivation of the non-local scaling in Supplement S1.3, we require that the distance between the triangles dist(*T, K*) ≥ *α* max(*h*_*T*_, *h*_*k*_) for some *α* > 0. By approximating the distance between two triangles as the distance between their center points, then this bound will certainly hold even in the cases where *T* and *K* are neighbors, justifying the summation over all triangles *K* ≠ *T* . Perhaps more concerning, however, is the calculation of this non-local term for every triangle *T*, as this summation indeed possesses significant computational memory and time requirements. To ease this computational burden, we take a simpler approach of only computing the nonlocal term for triangles within 3mm of *T*.

### 2.2 Electrode Preconditioner

The charge-based BEM-FMM formulation yields a well-conditioned linear system of second-kind Fredholm integral equations. However, on the voltage electrode-scalp interface, we also have a system of first-kind Fredholm integral equations corresponding to the Dirichlet boundary condition of the single-layer potential [18]. This Dirichlet constraint is a first-kind Fredholm integral equation. It also leads to physical and numerical singularities in the charge density and normal electric field along the electrode ring. These numerical issues are typically alleviated through the use of *N*_*e*_ × *N*_*e*_ preconditioner matrix *M*, where *N*_*e*_ is the number of triangles representing the electrodes on the scalp. This preconditioner is the discrete single-layer potential operator restricted to the electrode, with additional near-field corrections. The preconditioner serves the purpose of enforcing the Dirichlet boundary conditions on the electrode-scalp interface in a numerically stable manner [19–21].

During AMR procedure, the number of electrode triangles *N*_*e*_ grows substantially because the first-kind integral equation associated with the Dirichlet condition produces large numerical charge singularities near electrode boundaries (and thus the mesh is highly refined on and near the electrodes). Consequently, the resulting preconditioner will be computationally expensive/infeasible to build. Therefore, we opt to construct *M* in a block-wise fashion. This is done by sub-sampling the electrode faces evenly into a desired number of sectors *d* and constructing the preconditioner for each of these sectors.

The electrode sectors are determined in the following way. First, we locate the triangle nearest to the center of the electrode. From this triangle, we construct a local tangent plane with origin at the triangle center point. We emit *d* evenly spaced rays from the origin, creating *d* sectors of equal size. Each of the triangle center points across the electrode are then projected into this plane and labeled as belonging to the sector they land in. This procedure yields an approximately even distribution of triangles across sectors, enabling the construction of smaller, independent preconditioner blocks. An example of a sectored electrode is shown in Figure 1.

**Fig. 1:**
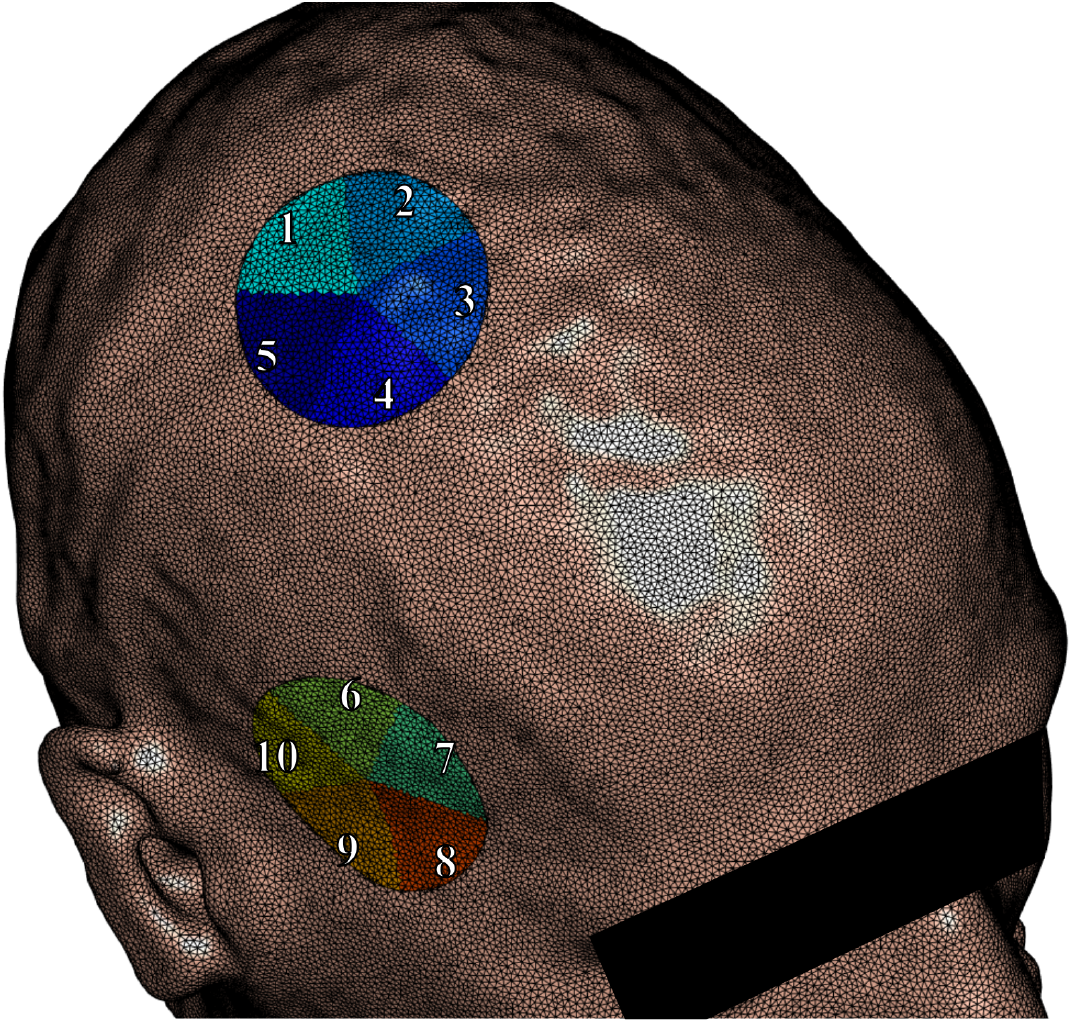
Example of electrodes split into five approximately equal sized sectors on the skin tissue surface. The preconditioner matrix *M* is constructed block-wise using each of these sectors.

### 2.3 Data Acquisition

For the head model segmentations described in the following section, we analyze 5 subject head models: two patients with drug-resistant major depressive disorder (N01 and N02), and three healthy participants (P01, P02, and P03), whose T1-weighted MRI data were provided to us by the authors of [22] and [23], respectively. Ethical approval of the study of [22] was obtained from the Human Research Protections Office at the University of New Mexico (UNM). The research was conducted in full compliance with the ethical standards outlined in the Declaration of Helsinki. Patients were recruited from the UNM Mental Health Center’s inpatient and outpatient services. All patients either had the decisional capacity to consent or, where necessary, provided assent with a surrogate decision-maker giving formal consent. Ethical approval of the study of [23] was obtained from the Ethics Committee of the Medical Faculty of Friedrich-Schiller-University Jena (Ethics Approval Number 2021-2347_2-BO). This study was also conducted in accordance with the Declaration of Helsinki and all volunteers provided written informed consent for participation and data use.

### 2.4 Head Models

We evaluate our adaptive BEM–FMM approach across three model types: the 5-layer sphere model, the 7-tissue SimNIBS headreco segmentation, and the 40-tissue Sim4Life head40 segmentation. These models span increasing levels of anatomical complexity, allowing us to assess the method’s behavior under conditions of varying realism. Conductivities for each model type and tissue are listed in Supplement S3. For each subject, we apply fixed voltages of ±1V at each voltage electrode. For subjects N01 and N02, we use electrode radii of 25mm; for subjects P01, P02, and P03, we consider electrode radii of 10mm and 5mm. For subjects P01, we also consider current electrodes (sponges) of 5mm radii with fixed currents of ±1mA.

#### 5-layer sphere

We use a benchmark case of 5 concentric spheres *S*_*i*_ of radii 92mm, 86mm, 80mm, 78mm,and 75mm. Two circular electrodes of radius 20mm are placed symmetrically on the outermost sphere *S*_1_. To provide a controlled deep-tissue target for TES field evaluation, we embed a pair of “hippocampi” modeled as the top and bottom hemispheres (relative to the electrode position) of a sphere of radius 50mm centered at the origin. This model structure is visualized in Figure 2.

**Fig. 2:**
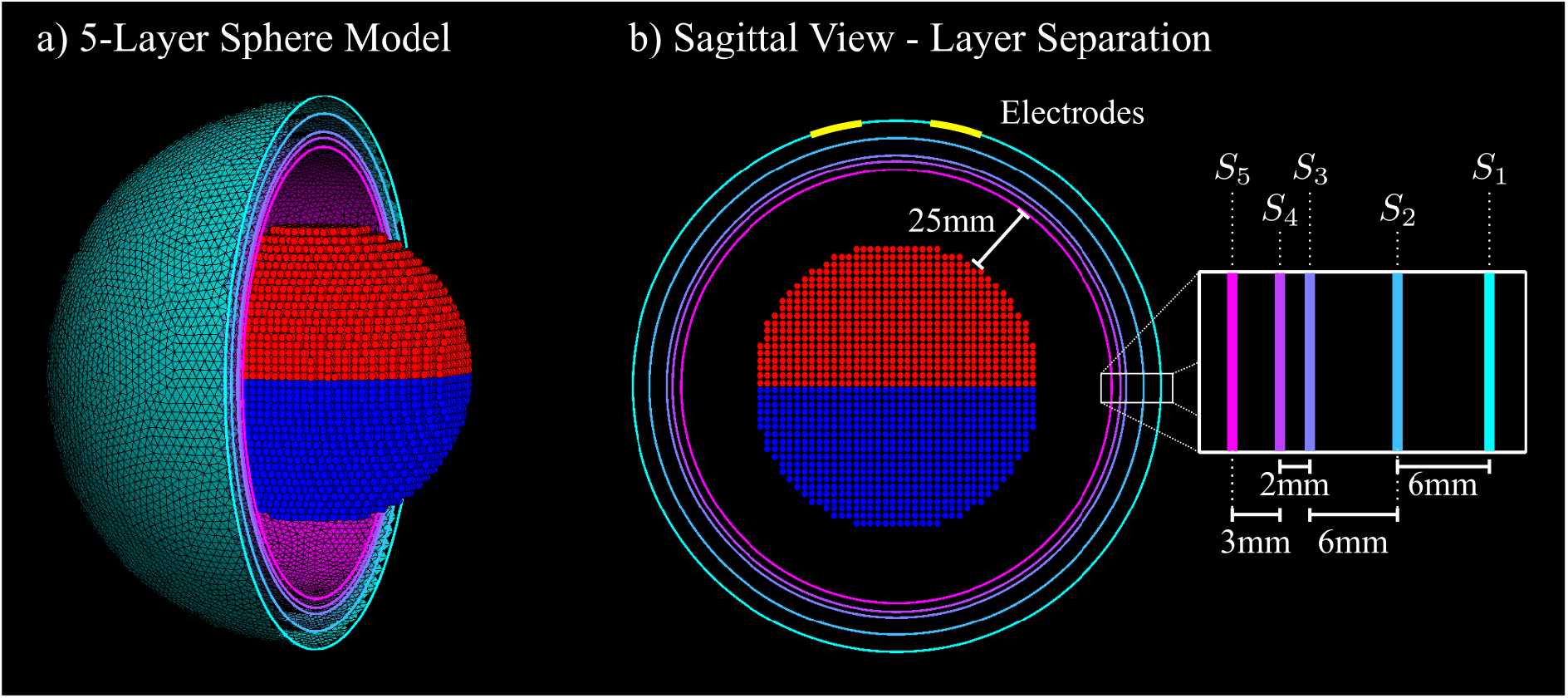
a) Visual of the 5-layer sphere model, with layers *Si* shown for *x* ≤ 0. The hemispheric grid of points representing the deep structures are shown as red and blue points. b) Sagittal view of the *x* = 0 cross-section, along with the separation distances of each layer as well as the distance from the inner-most layer to the deep structures. The position of the 20mm electrodes are depicted in yellow.

#### 7-tissue SimNIBS segmentation

The first head model type we use is the 7-tissue head model segmented via the SimNIBS headreco pipeline [15]. An example of the SimNIBS tissue segmentation is shown in Fig. 3a. Prior to refinement, the surface meshes are globally smoothed via non-manifold Taubin smoothing [24] and uniformly refined to remove segmentation artifacts and ensure geometric consistency across tissue interfaces.

**Fig. 3:**
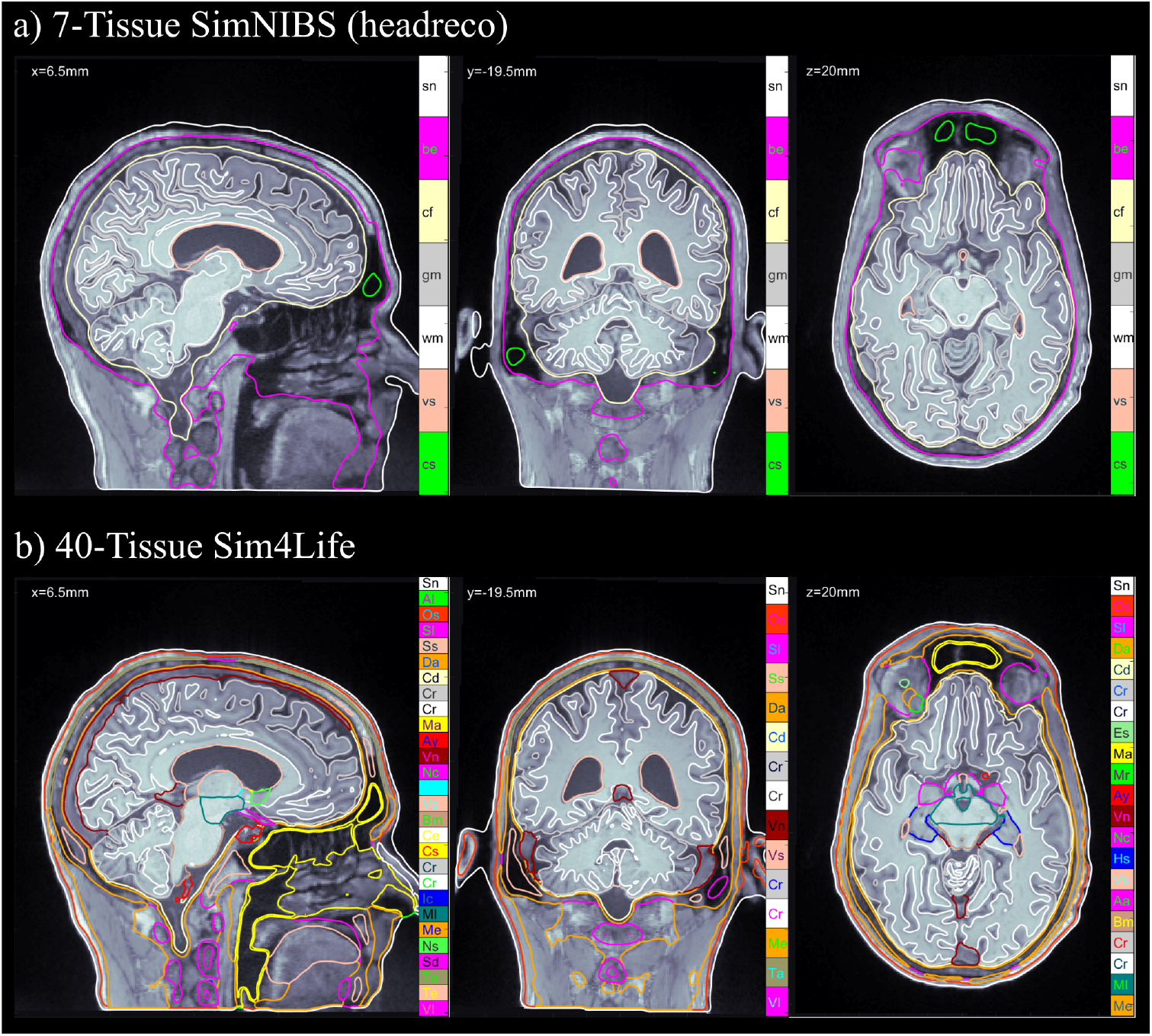
Subject N01 MRI volumes and overlayed tissue cross-sections of a) the 7-tissue SimNIBS headreco segmentation and b) the 40-tissue Sim4Life head40 segmentation in the sagittal (left), coronal (middle) and axial (right) planes. The full list of tissues and conductivities may be found in Supplement S3.

#### 40-tissue Sim4Life segmentation

The second head model type we use is the 40-tissue head model segmented via the IT’IS Foundation Sim4Life head40 segmentation [16, 25]. This segmentation includes detailed soft-tissue structures, thin layers, and non-manifold interfaces. This model represents the most challenging and fruitful scenario, with highly realistic, complex anatomical detail and conductivity contrasts. An example of the Sim4Life tissue segmentation is shown in Fig. 3b. As with the SimNIBS model, we apply initial global smoothing and uniform refinement. Global refinement is even more important in this case, where the tissue layers are very thin, since for accurate computation we should have the triangle edge lengths smaller than the distance between tissue boundaries. We note that some of the Sim4Life tissue surfaces have overlapping or shared triangles. In this case, the barycentric refinement can create non-manifold meshes with duplicate triangles or edges shared by three or more triangles. After refinement, we fix the mesh by removing these duplicated features.

### 2.5 Convergence Analysis

To assess the stability and reliability of our numerical models, we examine the convergence of three quantities of interest: the electrode currents *I*_*e*_, the normal electric field on the white matter surface **n** · **E**_WM_, and the magnitude of the volumetric electric field within the deep cortical structures (the hippocampi) **E**_HP_. These quantities are crucial for TES dose characterization as well as accurate EEG forward modeling. Particularly, stabilization of the electrode currents is essential for ensuring reliable numerical behavior, as they are highly sensitive to surface discretization. The total current on each electrode is computed by summing over the normal current flux through each of the electrode triangles.

For the head models, the hippocampal structures are obtained from the Sim4Life segmentation. We generate a set of interior grid points by applying MATLAB’s inpolyhedron function; these hippocampal points are reused for the SimNIBS model. These points serve as direct targets for volumetric field calculation at the end of each AMR step.

For these quantities, we compute the relative residual percentage

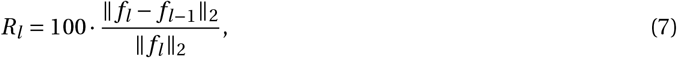

where *f*_*l*_ is the quantity of interest at the *l* -th AMR step. For the comparison of the normal electric field across iterations, we project the fields (which are evaluated on meshes refined via AMR) onto a reference mesh comprised of the white matter surface on the zeroth iteration (the initial mesh). The projections are computed by averaging the field values from the refined mesh over the *k*-nearest-neighbors of the reference mesh. To facilitate comparisons across subjects and segmentations, we will plot these convergence quantities against the model size relative to the initial model size, that is, the ratio of the total number of triangles at the each AMR step with respect to that of the initial model. For instance, the first step always has a relative model size of 4 since we always perform a uniform barycentric refinement on this step.

## 3 Results and Discussion

Fig. 4 shows the convergence results for the 5-layer sphere model with 20mm voltage electrodes. Fig. 5 shows the white matter and hippocampi relative percent errors, plotted against the relative model size, for subjects N01 and N02 with 25mm voltage electrodes. Fig. 6 shows the white matter relative percent errors, again plotted against the relative model size, for subjects P01, P02, and P03 using 10mm and 5mm voltage and current electrodes. Tables 1 and 2 show the convergence results for the 25mm and 10mm voltage electrodes, respectively. Table 3 shows the convergence results for the 5mm voltage and current electrodes. Figs. 7, 8, and 9 show the volumetric electric fields for subject P01 using the 10mm voltage and 5mm voltage and current electrodes. Note that in these results, the number of AMR iterations and the final model sizes can significantly vary. This is due to the inconsistent memory constraints of the servers upon which these simulations were run.

**Fig. 4:**
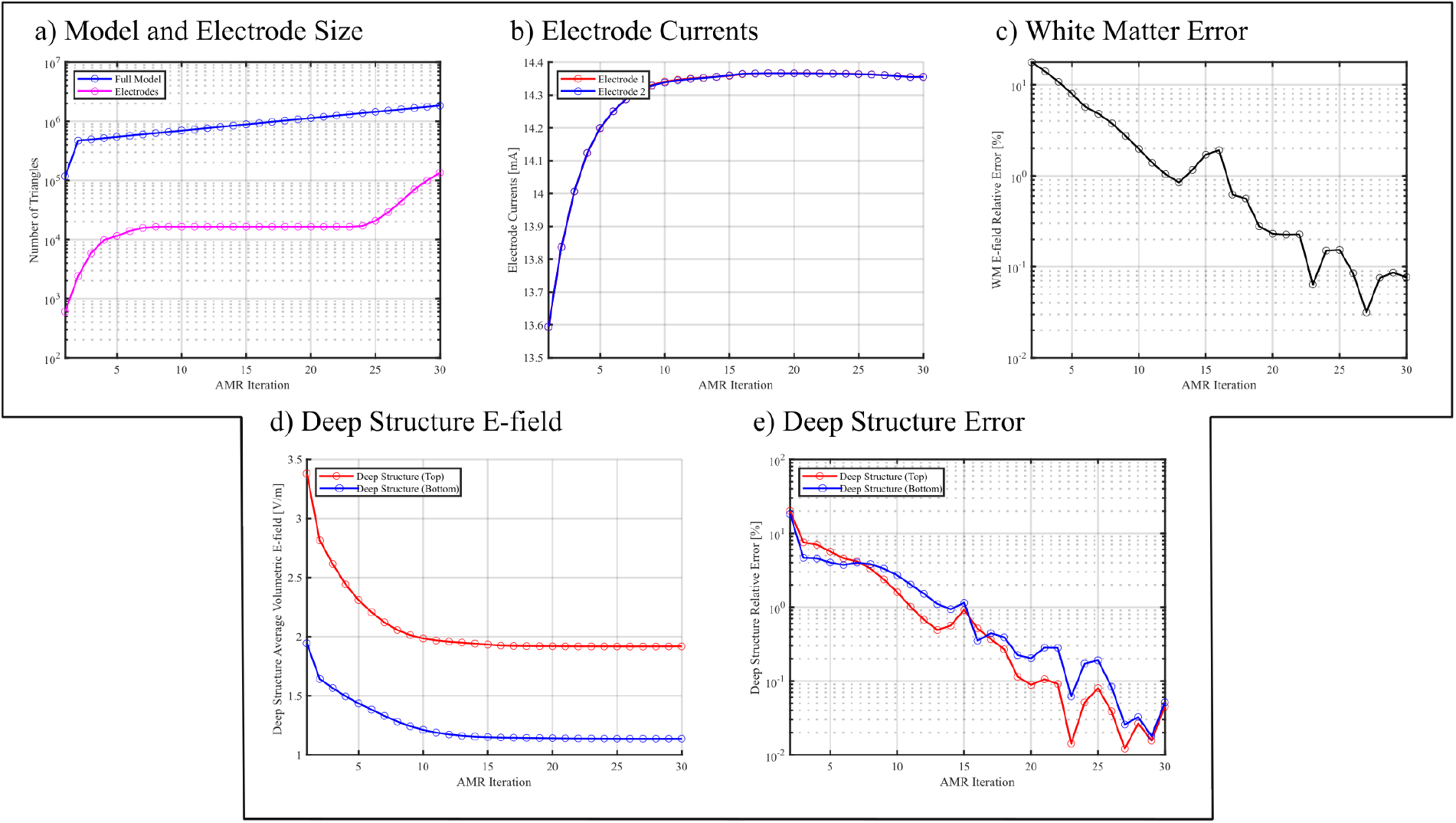
The convergence results vs. AMR iteration for the 5-layer sphere model depicted in Fig. 2 with 20mm voltage electrodes. a) The number of triangles used in the entire model and on the electrodes. b) The total current averaged over each electrode. c) The relative percent error of electric field on the “white matter” surface (in this case, the inner most layer *S*5). d) The average volumetric electric field of the deep structures depicted in Fig. 2. e) The relative percent error of the volumetric electric field on the deep structures.

**Fig. 5:**
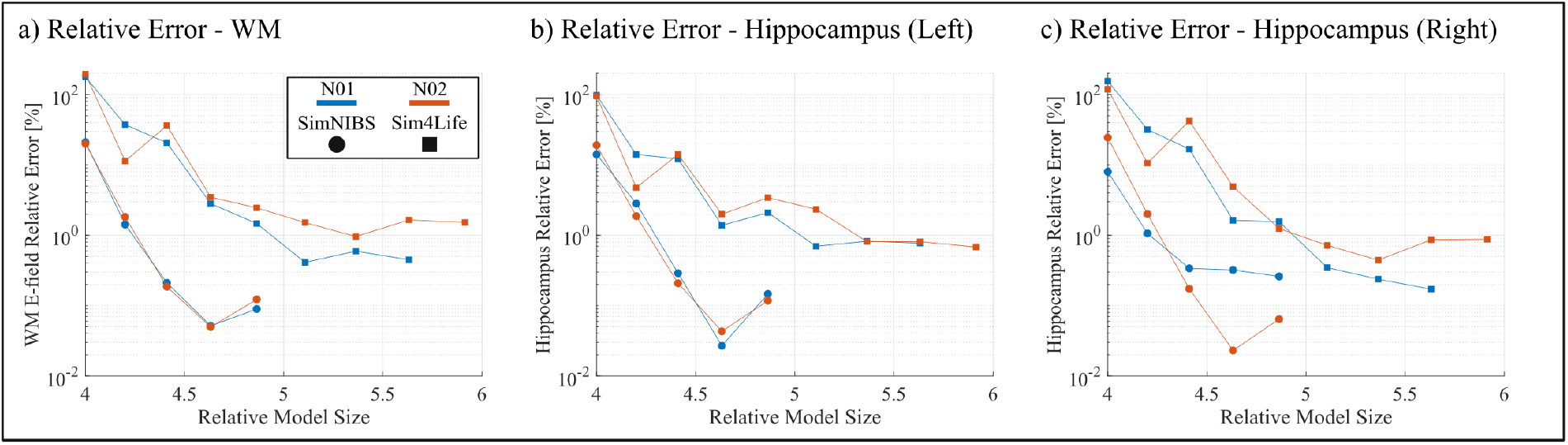
The convergence results for Subjects N01 and N02 with large voltage electrodes (25mm radius). a) The relative percent error of electric field on the white matter surface. b/c) The relative percent error of volumetric electric field on the left/right hippocampus. Subjects are delineated by color and model segmentations are delineated by shape. Errors are plotted against the number of triangles at each AMR step divided by that of the initial model (e.g. the initial error is always plotted at 4 because the first step is a uniform barycentric refinement).

**Fig. 6:**
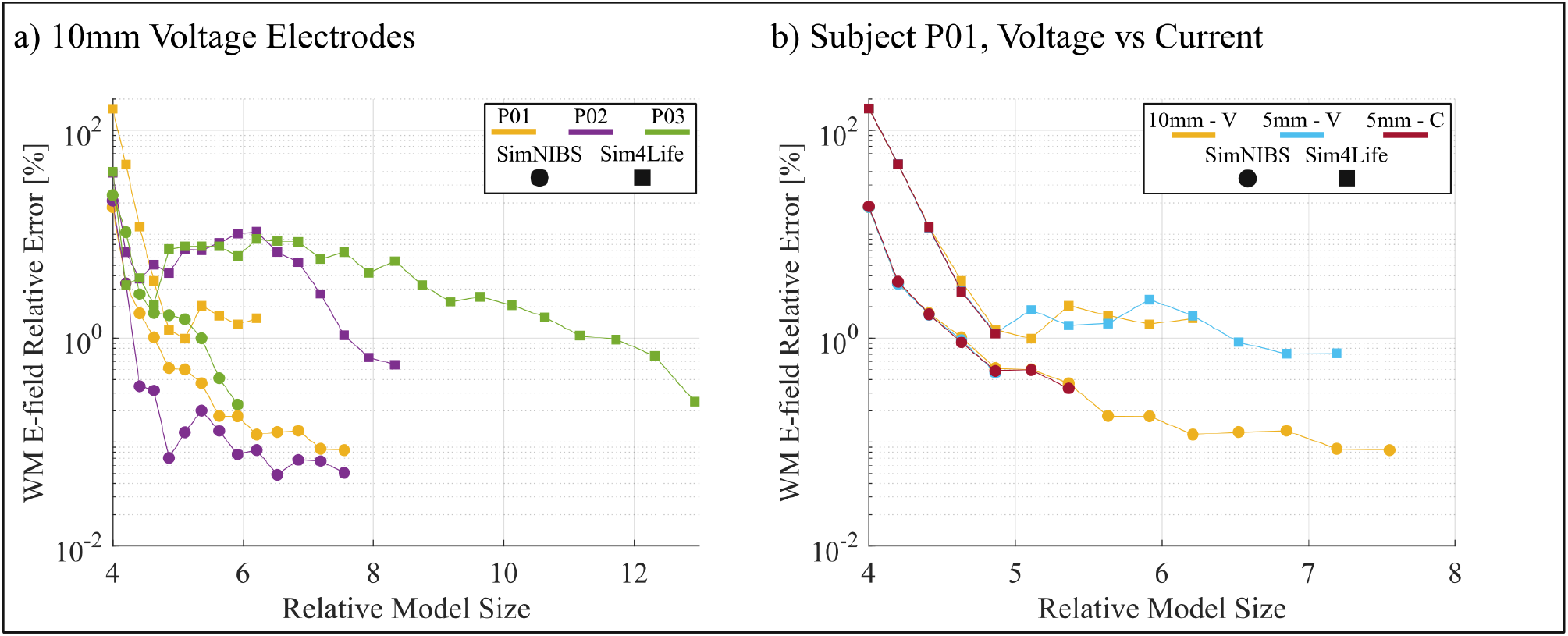
The convergence results for Subjects P01, P02, and P03 with small electrodes (10mm and 5mm radius).a) The relative percent error of electric field on the white matter surface using 10mm voltage electrodes. b) The relative percent error of electric field on the white matter surface on the subject P01 head model, using 10mm voltage electrodes, and 5mm voltage and current electrodes. Subjects (and electrode type) are delineated by color and model segmentations are delineated by shape. Errors are plotted against the number of triangles at each AMR step divided by that of the initial model (e.g. the initial error is always plotted at 4 because the first step is a uniform barycentric refinement).

**Table 1:** The convergence results for the models with 25mm voltage electrodes at the final AMR iteration. Columns from left to right indicate the model, final AMR iteration, total number of triangles, average electrode currents, relative percent error of electric field on the white matter, and the mean volumetric electric field on the right and left hippocampi.

| Model | Final Iteration | # Triangles [millions] | Current [mA] | WM % | HP (R) [V/m] | HP (L) [V/m] |
| --- | --- | --- | --- | --- | --- | --- |
| N01 (SimNIBS) | 6 | 19.4 | 23.00 | 0.089 | 2.493 | 1.483 |
| N01 (Sim4Life) | 9 | 25.0 | 14.68 | 0.448 | 0.960 | 0.679 |
| N02 (SimNIBS) | 9 | 16.8 | 21.78 | 0.122 | 1.870 | 1.284 |
| N02 (Sim4Life) | 10 | 23.5 | 15.39 | 1.516 | 0.890 | 0.743 |

**Table 2:**
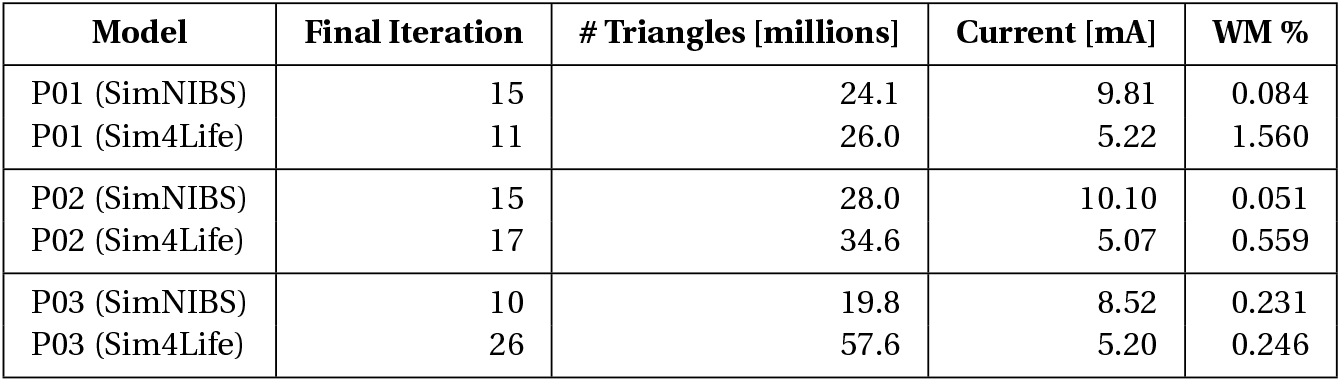
The convergence results for the models with 10mm voltage electrodes at the final AMR iteration. Columns from left to right indicate the model, final AMR iteration, total number of triangles, average electrode currents, and relative percent error of electric field on the white matter.

| Model | Final Iteration | # Triangles [millions] | Current [mA] | WM % |
| --- | --- | --- | --- | --- |
| P01 (SimNIBS) | 15 | 24.1 | 9.81 | 0.084 |
| P01 (Sim4Life) | 11 | 26.0 | 5.22 | 1.560 |
| P02 (SimNIBS) | 15 | 28.0 | 10.10 | 0.051 |
| P02 (Sim4Life) | 17 | 34.6 | 5.07 | 0.559 |
| P03 (SimNIBS) | 10 | 19.8 | 8.52 | 0.231 |
| P03 (Sim4Life) | 26 | 57.6 | 5.20 | 0.246 |

**Table 3:** The convergence results for the models with 5mm voltage and current electrodes at the final AMR iteration. Columns from left to right indicate the model, final AMR iteration, total number of triangles, average electrode currents or voltages, electrode impedance, and relative percent error of electric field on the white matter.

| Voltage Electrodes |  |  |  |  |  |
| --- | --- | --- | --- | --- | --- |
| Model | Final Iteration | # Triangles [millions] | Current [mA] | DC Resistance [$\Omega$] | WM % |
| P01 (SimNIBS) | 6 | 15.5 | 5.89 | 340 | 0.467 |
| P01 (Sim4Life) | 14 | 30.0 | 2.81 | 712 | 0.718 |

| Current Electrodes (Sponges) |  |  |  |  |  |
| --- | --- | --- | --- | --- | --- |
| Model | Final Iteration | # Triangles [millions] | Voltage [V] | DC Resistance [$\Omega$] | WM % |
| P01 (SimNIBS) | 8 | 17.1 | 0.3605 | 361 | 0.331 |
| P01 (Sim4Life) | 6 | 20.3 | 0.759 | 759 | 1.105 |

**Fig. 7:**
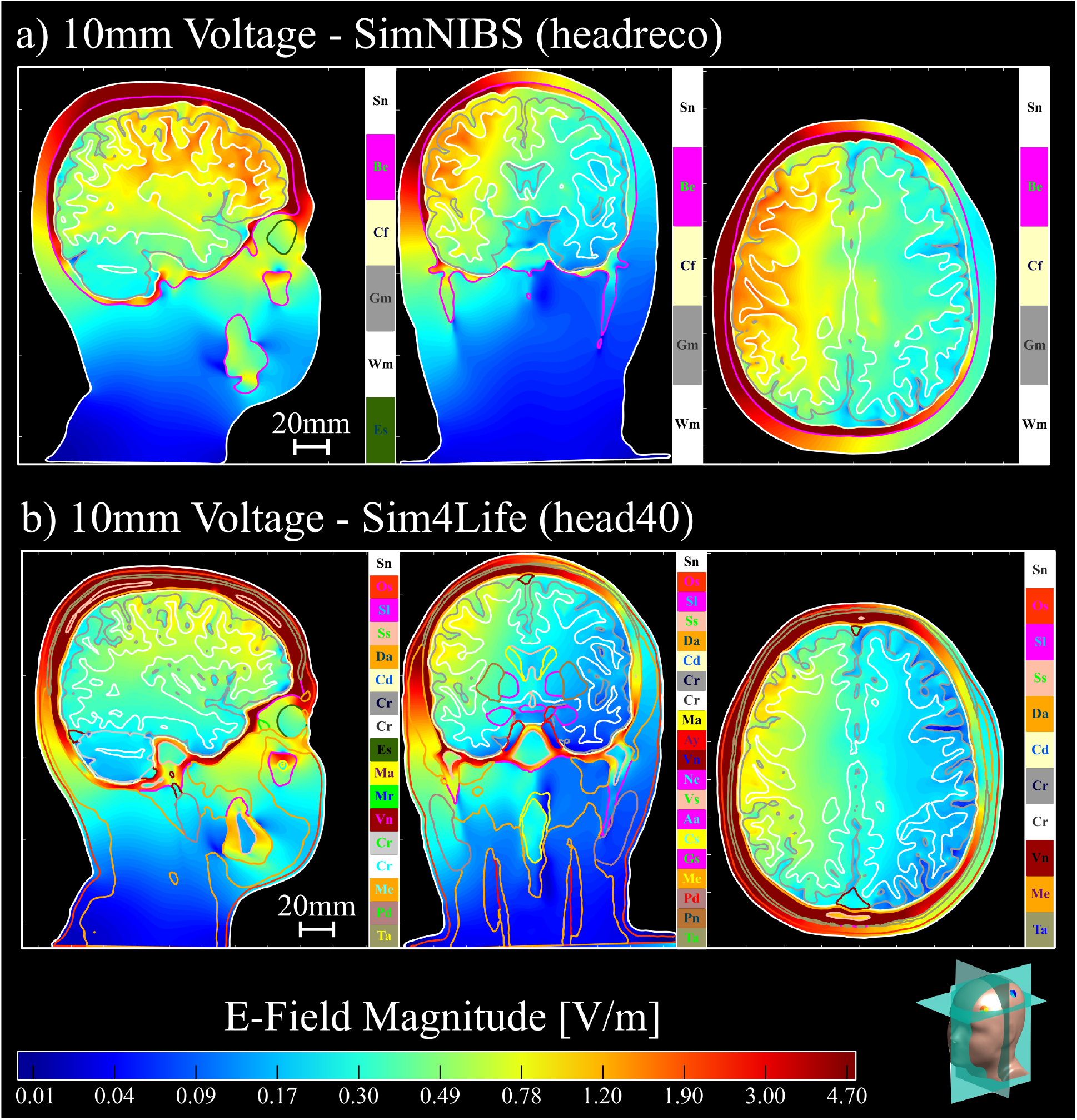
The volumetric electric fields of subject P01 for 10mm voltage electrodes with the a) SimNIBS (headreco) and b) Sim4Life (head40) segmentations. The electric field shown for the SimNIBS segmentation in a) are normalized by the currents shown in Tables 2 with respect to the Sim4Life segmentation. Electric fields are plotted on a logarithmic scale. Head model slices are visualized in the bottom right corner.

**Fig. 8:**
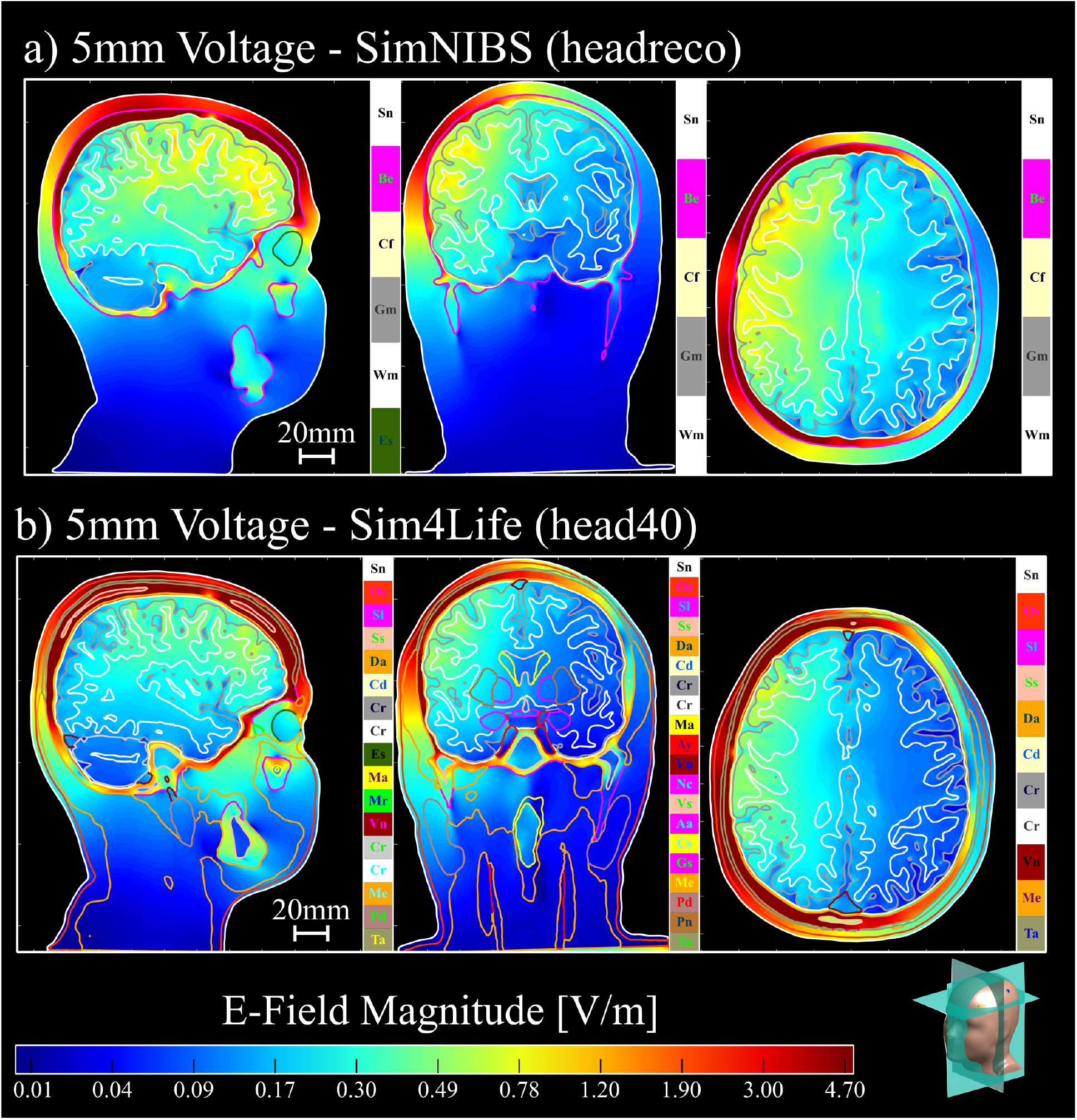
The volumetric electric fields of subject P01 for 5mm voltage electrodes with the a) SimNIBS (headreco) and b) Sim4Life (head40) segmentations. The electric field shown for the SimNIBS segmentation in a) are normalized by the currents shown in Tables 3 with respect to the Sim4Life segmentation. Electric fields are plotted on a logarithmic scale. Head model slices are visualized in the bottom right corner.

**Fig. 9:**
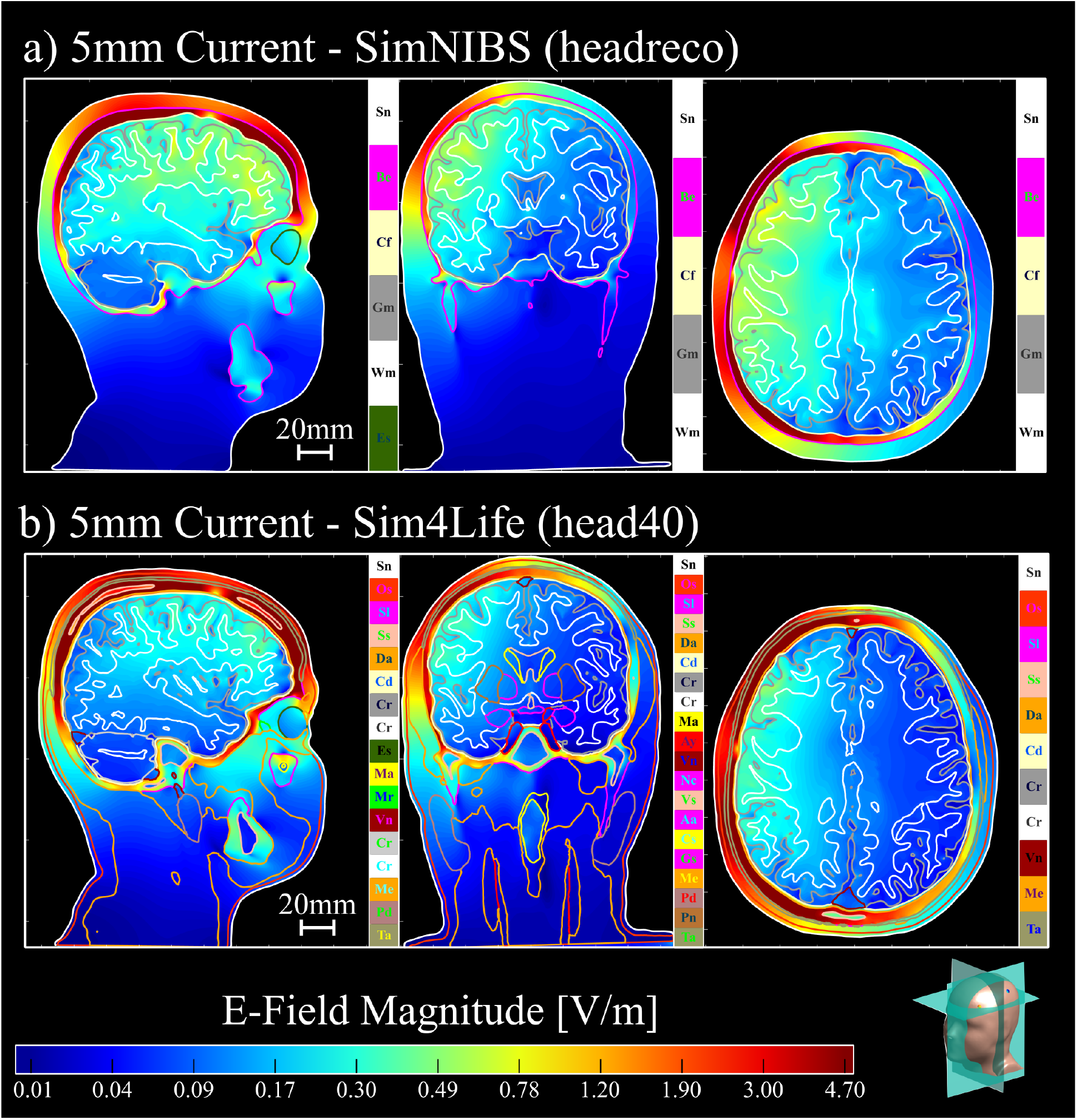
The volumetric electric fields of subject P01 for 5mm current electrodes with the a) SimNIBS (headreco) and b) Sim4Life (head40) segmentations. The electric field shown for the SimNIBS segmentation in a) are normalized by the currents shown in Tables 3 with respect to the Sim4Life segmentation. Electric fields are plotted on a logarithmic scale. Head model slices are visualized in the bottom right corner.

Fig. 4, serving as a baseline model, indicates that the refinement criterion of Equation 6 works well for ensuring convergence of the electric fields on the deepest target structures. We see a steady convergence of the electrode currents and deep structure electric fields in Figs. 4b and 4d, respectively. We see that when the model size reaches approximately 2 million triangles, we achieve relative percent errors below 0.1% for the “white matter” and deep structure measures in Figs. 4c and 4e. We also notice small “spikes” in the relative percent error measures: these may occur due to the accumulation of small numerical errors in the preconditioner calculation or FMM solver. These small errors are likely negligible as the model size increases, due to the overall steady reduction of the relative percent errors. These observations suggest that the charge residual refinement criterion serves as an effective criterion for determining convergent solutions.

In Fig. 5, we see that the behavior of the AMR solutions remains consistent across subjects N01 and N02 with the 25mm. For the SimNIBS segmentation, both subjects achieve a relative error near 0.1% at approximately the same rate for both the white matter and the left hippocampus in Fig. 5a and Fig. 5b, respectively. We see a slight deviation of this pattern in Fig. 5c, where subject N01 has slightly increased error over subject N02, but is sill below 0.2% error. For the Sim4Life segmentation, we see an increase in the error to slightly below 1%, but both subjects posses similar reduction rates across all target structures. This suggests that, for the large electrodes and fixed segmentation pipeline, the refinement criterion is fairly robust to anatomical variability among subjects.

For the 10mm voltage electrodes in Fig. 6a, the SimNIBS models for subjects P01, P02, and P03 all reach approximately 0.1% white matter relative error after a roughly 6-times increase in model size. By contrast, the Sim4Life models converge more slowly and with greater inter-subject variability, but each with each achieving an error near 1% at various refinement levels. This difference across segmentations is consistent with the increased geometric detail of the realistic Sim4Life tissue models; resolving the errors in charge solution on these more complex models indeed requires increased levels of mesh refinement. The results for P01 with 10mm and 5mm electrodes in Fig. 6b further show that reducing the electrode size does not fundamentally alter the behavior of the error rates for each segmentation. We see that the SimNIBS model reaches error near 0.4% with a roughly 5-times increase in model size, while the Sim4Life models achieve 1% error for a similar increase in model size.

In Tables 1 and 2, we see two additional trends. First, for voltage electrodes, the SimNIBS models consistently overestimate the electrode current when compared to the Sim4Life models. Second, for the 25mm electrodes, the mean hippocampal electric fields are lower in the Sim4Life models. This is likely due to the presence of many tissue layers in the Sim4Life segmentation which reduce field penetration deep within the cortex. (Note that the AMR procedure would not eliminate these differences since they arise from anatomical factors as opposed to numerical ones. Instead, the AMR ensures that the achieved electrode currents and electric fields converge with respect to the discretization, enabling these anatomical discrepancies to appear.)

Figs. 7, 8, and 9 illustrate these trends, where the electric fields for the SimNIBS segmentations show stronger field penetration into the cortical layers. We see that these field strengths are weaker when using the smaller electrodes in Fig. 8 and 9, as expected.

In Fig. 6b, we see that the behavior of the 5mm electrode solutions remains consistent for voltage and current electrodes, with almost no visible difference between the two for each segmentation. This is likely because the electrodes are sufficiently small enough so that the initial global refinement resolves the dominant component of the singularities at the electrode boundaries. In Table 3, we see that the achieved error for both voltage and current electrodes is comparable relative to the final model size. When normalizing the voltage electrode results by current, we see that the effective electrode impedance for the SimNIBS model is lower than that of the Sim4Life model, and this effect is maintained for the current electrodes. This is intuitive, again due to the complex tissue structure of the Sim4Life model. The difference in total impedance when comparing voltage and current electrodes is likely due to small numerical errors or differences in total number of adaptive passes. This impedance difference leads to the differences in electric field strengths seen when comparing Figs. 8a and 8b to Figs. 9a and 9b, respectively, where the fields from current electrodes are noticeably weaker in the deeper cortical regions.

## 4 Conclusions

In this work, we have established an adaptive mesh refinement strategy for the charge based BEM-FMM formulation of TES and EEG forward modeling. We have derived an error estimator which captures both local and nonlocal contributions of the single-layer potential operator. From this, we constructed a surrogate refinement criterion based on the difference in charge solutions from each AMR step. We analyzed the performance of this criterion on three model types through the relative residual error of the white matter and deep target electric fields. Our method was validated for a 5-layer sphere model, with deep cortical targets yielding relative errors below 0.1%. Performance on head models yielded relative errors near 0.1% for the 7-tissue SimNIBS models and 1% for the 40-tissue Sim4Life models across all subjects and electrode sizes. Overall we found that the SimNIBS segmentation led to an overestimation of the electrode currents and correspond electric field strengths compared to the Sim4Life models. Thorugh these results, we have demonstrated that our AMR criterion is effective at determining stable TES and EEG forward solutions in realistic head models.

## Acknowledgments

D.A.D, G.M.N, and S.N.M. were supported by the NIBIB Grant 1R01EB035484, and the NIMH Grant 1R01MH130490. H.O. received funding by the Carl Zeiss Stiftung under the CZS Breakthroughs program within the “PollenNet” project (Grant number: P2022-08-006). J.H. received funding from the German Federal Ministry of Education and Research (BMBF) grant DryPole (01GQ2304A) and the Free State of Thuringia (2018 IZN 004), co-financed by the European Union under the European Regional Development Fund (ERDF). Z.D. was supported by the National Institute of Mental Health Intramural Research Program (ZIAMH002955).

## Author Contributions

D.A.D wrote the main manuscript text. D.A.D., G.M.N., and S.N.M. developed the methodology and algorithms used to generate the results. H.O., J.H., and Z.D. collected, processed, and provided the data analyzed. All authors reviewed, edited, and revised the manuscript.

## Data Availability

The data and code may be made available upon request.

## Notes

### Competing Interest Statement

The authors have declared no competing interest.

### Summary of Updates

Misspelling in title, changed "Simulation" to "Stimulation"

## References

[1] Ren, C. et al. Transcranial electrical stimulation in treatment of depression: A systematic review and meta-analysis. JAMA Network Open 8, e2516459 (2025).

[2] Mukhtar, F., Regenold, W. & Lisanby, S. H. Recent advances in electroconvulsive therapy in clinical practice and research. Faculty Reviews 12 (2023).

[3] Chaddad, A., Wu, Y., Kateb, R. & Bouridane, A. Electroencephalography signal processing: A comprehensive review and analysis of methods and techniques. Sensors 23, 6434 (2023).

[4] Nuñez Ponasso, G. et al. Improving eeg forward modeling using high-resolution five-layer bem-fmm head models: Effect on source reconstruction accuracy. Bioengineering 11, 1071 (2024). URL https://www.mdpi.com/2306-5354/11/11/1071.

[5] Makarov, S. N. et al. Boundary element fast multipole method for enhanced modeling of neurophysiological recordings. IEEE Transactions on Biomedical Engineering 68, 308–318 (2021).

[6] Weise, K., Wartman, W. A., Knösche, T. R., Nummenmaa, A. R. & Makarov, S. N. The effect of meninges on the electric fields in tes and tms. numerical modeling with adaptive mesh refinement. Brain Stimulation: Basic, Translational, and Clinical Research in Neuromodulation 15, 654–663 (2022).

[7] Makaroff, S. N. et al. A fast direct solver for surface-based whole-head modeling of transcranial magnetic stimulation. Scientific Reports 13 (2023).

[8] Wartman, W. A. et al. Fast eeg/meg bem-based forward problem solution for high-resolution head models. NeuroImage 306, 120998 (2025). URL https://www.sciencedirect.com/science/article/pii/S1053811924004956.

[9] Babuvška, I. & Rheinboldt, W. C. Error estimates for adaptive finite element computations. SIAM Journal on Numerical Analysis 15, 736–754 (1978).

[10] Ainsworth, M. & Tinsley Oden, J. A posteriori error estimators for second order elliptic systems: Part 1. theoretical foundations and a posteriori error analysis. Computers & Mathematics with Applications 25, 101–113 (1993).

[11] Carstensen, C. & Stephan, E. P. Adaptive coupling of boundary elements and finite elements. ESAIM: Mathematical Modelling and Numerical Analysis 29, 779–817 (1995).

[12] Gantumur, T. Adaptive boundary element methods with convergence rates. Numerische Mathematik 124, 471–516 (2013).

[13] Feischl, M., Führer, T., Heuer, N., Karkulik, M. & Praetorius, D. Adaptive boundary element methods: A posteriori error estimators, adaptivity, convergence, and implementation. Archives of Computational Methods in Engineering 22, 309–389 (2014).

[14] Wartman, W. A. et al. An adaptive h-refinement method for the boundary element fast multipole method for quasi-static electromagnetic modeling. Phys. Med. Biol. 69, 055030 (2024).

[15] Simnibs standard conductivity values. https://simnibs.github.io/simnibs/build/html/documentation/conductivity.html (2025).

[16] Automated head segmentation in sim4life v8.0.1: A new reference. https://zmt.swiss/news-and-events/news/sim4life/automated-head-segmentation/ (2024).

[17] Makarov, S. N., Noetscher, G. M., Raij, T. & Nummenmaa, A. A quasi-static boundary element approach with fast multipole acceleration for high-resolution bioelectromagnetic models. IEEE Transactions on Biomedical Engineering 65, 2675–2683 (2018).

[18] Nuñez Ponasso, G. A survey on integral equations for bioelectric modeling. Physics in Medicine &; Biology 69, 17TR02 (2024). URL 10.1088/1361-6560/ad66a9.

[19] Steinbach, O. & Wendland, W. The construction of some efficient preconditioners in the boundary element method. Advances in Computational Mathematics 9, 191–216 (1998).

[20] Stevenson, R. & van Venetië, R. Uniform preconditioners for problems of negative order. Mathematics of Computation 89, 645–674 (2019).

[21] Giunzioni, V., Ortiz G. J. E., Merlini, A., Adrian, S. B. & Andriulli, F. P. On a calderón preconditioner for the symmetric formulation of the electroencephalography forward problem without barycentric refinements. Journal of Computational Physics 491, 112374 (2023).

[22] Dib, M., Lewine, J. D., Abbott, C. C. & Deng, Z.-D. Electroconvulsive therapy modulates loudness dependence of auditory evoked potentials: a pilot meg study. Frontiers in Psychiatry 15 (2024). URL https://www.frontiersin.org/journals/psychiatry/articles/10.3389/fpsyt.2024.1434434.

[23] Oppermann, H., Bruña, R., Güllmar, D., Klingner, C. & Haueisen, J. Characterization of transient events during intermittent photic stimulation. NeuroImage 323, 121587 (2025).

[24] Taubin, G. Curve and surface smoothing without shrinkage. In Proceedings of IEEE International Conference on Computer Vision, ICCV-95, 852–857 (IEEE Comput. Soc. Press, 1995).

[25] Sim4life tissue models. https://sim4life.swiss/#tissue-models (2025).

[26] Ding, Z. A proof of the trace theorem of sobolev spaces on lipschitz domains. Proceedings of the American Mathematical Society 124, 591–600 (1996).

[27] Sauter, S. Boundary Element Methods. SpringerLink (Springer-Verlag Berlin Heidelberg, Berlin, Heidelberg, 2011). Includes bibliographical references and indexes.

[28] Lax, P. D. & Milgram, A. N. IX. Parabolic Equations, 167–190 (Princeton University Press, 1955).

[29] Cea, J. Approximation variationnelle des problèmes aux limites. Annales de l’Institut Fourier 14, 345–444 (1964).

[30] Steinbach, O. (ed.) Numerical Approximation Methods for Elliptic Boundary Value Problems. Springer-Link (Springer Science+Business Media, LLC, New York, NY, 2008). Includes bibliographical references (p. [375]-382) and index.

[31] Brenner, S. C. & Scott, L. R. The Mathematical Theory of Finite Element Methods (Springer New York, 2008).

[32] Baumgartner, C. et al. IT’IS Database for thermal and electromagnetic parameters of biological tissues. https://itis.swiss/database (2025). xAccessed December 3, 2025.

