## Supplements for "Charge Based Boundary Element Method with Residual Driven Adaptive Mesh Refinement for High Resolution Electrical Stimulation Modeling"

April 2026

### S1 Supplement: Derivation of Refinement Criterion

#### S1.1 Galerkin Solution of the Single Layer Potential

We work in the context of a single layer potential

$$u(x) = \int_S \rho(y)\Phi(x,y)\,dS_y \tag{8}$$

where $\Phi(x,y) = 1/(4\pi|x-y|)$ is the fundamental solution to the Laplace equation in $\mathbb{R}^3$ and $S$ is the Lipschitz boundary of a compact domain $\Omega \subset \mathbb{R}^3$ (this boundary can be a union of disjoint Lipschitz boundaries $S = S_1 \cup S_2 \cup \cdots \cup S_n$). The single layer potential can be extended to a general operator,

$$\left(\mathscr{S}(\varphi)\right)(x) := \left[\int_S \varphi(x)\Phi(x,y)\,dS_y\right]\bigg|_S, \tag{9}$$

which restricts the potential to the boundary $S$. Here $\mathscr{S} : \mathscr{H}^{-1/2}(S) \to \mathscr{H}^{1/2}(S)$ where $\mathscr{H}^s$ is the Sobolev space of order $s$ with corresponding dual $\mathscr{H}^{-s}$. The single layer potential operator $\mathscr{S}$ is a bounded elliptic operator [13]. We can therefore define the energy norm

$$|||\varphi||| := \langle \mathscr{S}\varphi, \varphi\rangle^{1/2}_{L_2(S)} \tag{10}$$

for $\varphi \in \mathscr{H}^{-1/2}$ [12]. (In what follows, we drop the subscript on the inner product when clear).

Note that the charge density $\rho$ indeed lies in $H^{-1/2}(S)$. This may be seen by considering that the potential $u \in H^1(S)$ by definition. So the trace mapping $\gamma u = u|_S \in H^{1/2}(S)$ [26] and the normal derivatives $\partial_n u|_S \in H^{-1/2}$ [27]. From the jump relations [18]:

$$\partial_n u_\pm(x) = \int_S \rho(x) \frac{\partial \Phi(x,y)}{\partial n(x)} dS_y \mp \frac{1}{2}\rho(x), \tag{11}$$

we see that $\rho = \partial_n u_- - \partial_n u_+ \in H^{-1/2}(S)$.

Since $\rho \in \mathscr{H}^{-1/2}(S)$, for a fixed $f \in \mathscr{H}^{1/2}$ we can consider the operator equation

$$\mathscr{S}\rho = f. \tag{12}$$

For a closed linear subspace $\omega \subset \mathscr{H}^{-1/2}(S)$, the Galerkin approximation $\rho_h \in \omega$ of $\rho$ is characterized by

$$\langle \mathscr{S}\rho_h, v\rangle = \langle f, v\rangle \quad \forall v \in \mathscr{H}^{-1/2}. \tag{13}$$

The existence and uniqueness of solutions to 13 follows from the fact that $\mathscr{S}$ is a bounded elliptic operator and by the Lax-Milgram Theorem [28]. Furthermore, the existence of a "best" Galerkin solution follows from Céa's Lemma [29].

#### S1.2 Adaptive BEM Criteria

Denote by $\mathscr{T}$ the triangulation of the boundary $S$, and $\rho_\tau$ the Galerkin solution over the mesh $\tau \subseteq \mathscr{T}$. The Galerkin residual $r_\tau = f - \mathscr{S}\rho_\tau$ is often used as a starting point for determine local and global error estimates. However, in the context of charge-based BEM-FMM, the boundary data $f$ is a priori unknown, and thus the exact Galerkin residual is not calculable. This presents a problem in that the standard adaptive BEM criteria is not directly applicable here. Therefore, for our purposes, we seek an error estimator $\eta_\tau$ such that $\|r_\tau\| \le C\eta_\tau$ for some $C > 0$. Such an estimator is called *reliable*, since a decreasing $\eta_\tau$ in adaptive iterations guarantees a decreasing Galerkin residual.

We begin with the norm equivalence of the Galerkin residual

$$\alpha\|\rho - \rho_\tau\|_{\mathscr{H}^{-1/2}(S)} \le \|r_\tau\|_{\mathscr{H}^{1/2}(S)} \le \beta\|\rho - \rho_\tau\|_{\mathscr{H}^{-1/2}(S)} \tag{14}$$

for some constants $\alpha, \beta > 0$ [12]. (In the classical adaptive BEM theory, this equivalence is used to justify using the Galerkin residual as an appropriate estimator for the charge residual, the true quantity of interest. However, here we are interested in simultatneously bounding both residuals.) Denoting by $e_\tau = \rho - \rho_\tau$ as the charge residual, we have that

$$\begin{aligned} \|r_\tau\|_{\mathscr{H}^{1/2}(S)} &= \|f - \mathscr{S}\rho_\tau\|_{\mathscr{H}^{1/2}(S)}, \\ &= \|\mathscr{S}e_\tau\|_{\mathscr{H}^{1/2}(S)}, \\ &\le \|\mathscr{S}\|_{\text{op}}\|e_\tau\|_{\mathscr{H}^{-1/2}(S)}, \end{aligned} \tag{15}$$

where $\|\cdot\|_{\text{op}}$ is the natural operator norm.

The norm $\|\cdot\|_{\mathscr{H}^{-1/2}(S)}$ is not directly computable, but we possess the norm equivalence of the energy norm [12]

$$\alpha\|\cdot\|^2_{\mathscr{H}^{-1/2}(S)} \le |||\cdot||| \le \beta\|\cdot\|^2_{\mathscr{H}^{-1/2}(S)}, \tag{16}$$

which provides the charge residual bound

$$\begin{aligned} \|e_\tau\|_{\mathscr{H}^{-1/2}(S)} &\le c|||e_\tau|||^{1/2}, \\ &= \langle \mathscr{S}e_\tau, e_\tau\rangle^{1/2}, \end{aligned} \tag{17}$$

for some $c > 0$.

The bound given by Equation 17 is a global bound, i.e. a bound over all triangles $T \in \tau$. We can split this bound into two parts, one considering the local accumulation of charge error in a single triangle $T$, and the other taking into account the interaction of two triangles $T$ and $K$. First, we have the discrete formulation of the energy norm over a triangles $T, K \in \tau$,

$$\langle \mathscr{S} e_T, e_K \rangle = e_T^{\mathrm{T}} \tilde{\mathscr{S}}_{T,K} e_K, \tag{18}$$

where $\tilde{\mathscr{S}}_{T,K}$ are the entries of the linearization of $\mathscr{S}$ given by

$$\tilde{\mathscr{S}}_{T,K} = \int_T \int_K \Phi(x, y) dS_x dS_y. \tag{19}$$

In Section S1.3, we find the local and non-local scalings of $\tilde{\mathscr{S}}$:

$$\tilde{\mathscr{S}}_{T,T} = c_1 |T|^{3/2}, \tag{20}$$

and

$$\tilde{\mathscr{S}}_{T,K} = c_2 \frac{|T||K|}{\mathrm{dist}(T,K)}, \tag{21}$$

for $c_1, c_2 > 0$, where $|T|$ and $|K|$ denote the triangle areas, and $\mathrm{dist}(T,K) = \min_{x \in T, y \in K} |x - y|$. Therefore, we can split apart the discrete operator $\tilde{\mathscr{S}}_{T,K}$ in Equation 18, and write the global error estimator $\eta_\tau$ as

$$\langle \mathscr{S} e_\tau, e_\tau \rangle \approx \eta_\tau = c_1 \sum_{T \in \tau} \left( |T|^{3/2} e_T^{\mathrm{T}} e_T \right) + \frac{1}{2} c_2 \sum_{\substack{T,K \in \tau \\ T \neq K}} \left( \frac{|T||K|}{\mathrm{dist}(T,K)} e_T^{\mathrm{T}} e_K \right), \tag{22}$$

where the 1/2 factor in the non-local term is used to avoid double counting, since the discrete operator $\mathscr{S}_{T,K}$ is symmetric.

By combining Equations 15, 17, and 22, we find a global bound on the Galerkin residual,

$$\| r_\tau \|_{\mathscr{H}^{1/2}(S)} \leq \eta_\tau^{1/2}. \tag{23}$$

To utilize this in a practical way as a criterion for adaptive mesh refinement, we may consider the local error estimate as

$$\eta_T = c_1 |T|^{3/2} e_T^{\mathrm{T}} e_T + \frac{1}{2} c_2 \sum_{\substack{K \in \tau \\ T \neq K}} \left( \frac{|T||K|}{\mathrm{dist}(T,K)} e_T^{\mathrm{T}} e_K \right), \tag{24}$$

where now the non-local term is computed for the single triangle $T$. This provides us with a bound on the local residual

$$\| r_T \|_{\mathscr{H}^{1/2}(S)} \leq \eta_T^{1/2}. \tag{25}$$

Unfortunately, as the charge $\rho$ is unknown, the local and global charge residuals $e_T$ and $e_\tau$ are not computable. Therefore, we search for a sufficient choice of surrogate function $\xi_\tau \in \mathscr{H}^{-1/2}(S)$ to substitute for the charge residual $e_\tau$. By "sufficient choice," we mean that the *surrogate discrepancy*

$$D(\xi_\tau) = ||| e_\tau - \xi_\tau ||| \tag{26}$$

tends to zero in the limit of adaptive mesh refinement iterations. In particular, denoting by $l$ the current adaptive iteration with $\rho_l$ being the charge solution and $A_{l,T}$ the area of triangle $T$ at iteration $l$, we find that the (local) surrogate

$$\xi_{l,T} := A_{l,T}^p \left( \rho_l - \rho_{l-1} \right) \tag{27}$$

is sufficient for $p \geq 0$, see Section S1.4. In practice, we find that $p = 3/2$ tended to yield the best and most consistent results across models.

#### S1.3 Proof of Non-Local and Local Scalings

In what follows, we denote the distance between two triangles $T$ and $K$ by $d_{T,K} = \text{dist}(T,K)$. We also denote the diameter of $T$ by $h_T = \max_{x,x' \in T} |x - x'|$. We first prove the non-local scaling, which is relatively simple.

**Proposition S1.1.** *For two triangles $T, K \in \tau$ such that $d_{T,K} \geq \alpha \max(h_T, h_K)$ for some $\alpha > 0$, we have the non-local scaling $\tilde{\mathscr{S}}_{T,K} = c\frac{|T||K|}{d_{T,K}}$ for some $c > 0$.*

*Proof.* For any $x \in T$ and $y \in K$, $|x - y| \leq d_{T,K}$. So we have the upper bound

$$\frac{1}{|x-y|} \leq \frac{1}{d_{T,K}}. \tag{28}$$

Now take $x_0, \in T$ and $y_0 \in K$ such that $|x_0 - y_0| = d_{T,K}$. Then by the triangle inequality

$$\begin{aligned} |x-y| &\leq |x - x_0| + |x_0 - y_0| + |y - y_0|, \\ &= h_T + d_{T,K} + h_K, \\ &\leq (1 + \frac{2}{\alpha}) d_{T,K}. \end{aligned} \tag{29}$$

Therefore, we have the lower bound

$$\frac{1}{(1+\frac{2}{\alpha})d_{T,K}} \leq |x-y|. \tag{30}$$

Integrating the inequality chain over $T$ and $K$, we find

$$\frac{|T||K|}{(1+\frac{2}{\alpha})d_{T,K}} \leq 4\pi \int_T \int_K \frac{1}{4\pi|x-y|} dS_x dS_y \leq \frac{|T||K|}{d_{T,K}}, \tag{31}$$

or, rather,

$$c_1 \frac{|T||K|}{d_{T,K}} \leq \tilde{\mathscr{S}}_{T,K} \leq c_2 \frac{|T||K|}{d_{T,K}}. \tag{32}$$

□

Note that the inequality $d_{T,K} \geq \alpha \max(h_T, h_K)$ is very easy to check for any given mesh. Assuming the mesh $\tau$ is shape regular, then choosing the parameter $\alpha$ should be straight-forward. For example, if we approximate the dist$(T,K)$ as the distance between the center points of the triangles, then there is certainly $\alpha > 0$ which satisfies the condition of Proposition S1.1. A natural question then arises: should $\alpha$ be fixed, or can it change for every adaptive pass? This is certainly worth further exploration.

Before proving the local scaling, we need some preliminaries on the mesh. Let $x_1, x_2, x_3$ be the nodes of triangle $T$, with pairwise edges $e_1, e_2, e_3$. Denote by $\hat{T}$ the reference triangle corresponding to the mesh $\tau$. We have a change of coordinates $F_T : \hat{T} \to T$ from the reference triangle to the the mesh, defined by the affine map [30]

$$F_T(\hat{x}) = B_T \hat{x} + x_1, \tag{33}$$

where

$$B_T = \begin{bmatrix} (x_2 - x_1)_1 & (x_3 - x_2)_1 \\ (x_2 - x_1)_2 & (x_3 - x_2)_2 \\ (x_2 - x_1)_3 & (x_3 - x_2)_3 \end{bmatrix} = [e_1 \quad e_2] \tag{34}$$

Taking $\xi_1, \xi_2$ as a local parameterization of $T$, we can write the Jacobian as the linear mapping $dF_T : \hat{T} \to T$:

$$dF_T(\hat{x}) = \begin{bmatrix} \partial_{\xi_1} F_T & \partial_{\xi_2} F_T \end{bmatrix} \hat{x} \tag{35}$$

**Lemma S1.2.** *The Jacobian possesses the scaling* $\|dF_T\| = ch_T$ *for some* $c > 0$.

*Proof.* Taking the operator norm as the longest column vector of $dF_T$, and since the tangent vectors $\partial_{\xi_i} F_T$ lie in the local tangent plane of $T$, we have for some $c_1 > 0$

$$\|dF_T\| \leq c_1 h_T. \tag{36}$$

Conversely, each edge $e_i = B_T \hat{e}_i$ for edge $\hat{e}_i$ on the reference triangle, with $\|\hat{e}_i\| = 1$. Then we have

$$\begin{aligned} h_T &= \max_i \|e_i\|, \\ &\leq \|e_i\|, \\ &= \|B_T \hat{e}_i\|, \\ &\leq \|B_T\| \|\hat{e}_i\|, \\ &\leq \|dF_T\|. \end{aligned} \tag{37}$$

That is, we have

$$h_T \leq \|dF_T\| \leq c_1 h_T. \tag{38}$$

□

**Lemma S1.3.** *The singular values of the Jacobian satisfy* $\sigma_{\min} = c_1 h_T$ *and* $\sigma_{\max} = c_2 h_T$ *for some* $c_1, c_2 > 0$.

*Proof.* By definition of the operator norm, $\|dF_T\| = \sigma_{\max}$. So $\sigma_{\max} = c_1 h_T$ for some $c_1 > 0$ by Lemma S1.2. Further,

$$\sigma_{\min}\sigma_{\max} = \|\partial_{\xi_1} F_T \times \partial_{\xi_2} F\| = \|\partial_{\xi_1} F_T\| \|\partial_{\xi_2} F_T\| = c_2 h_T^2, \tag{39}$$

where the last equality holds again by Lemma S1.2. Thus, $\sigma_{\min} = c_2 h_T$ for some $c_2 > 0$. □

We now have what we need to prove the local scaling.

**Proposition S1.4.** *For each triangle* $T \in \tau$, *we have the local scaling* $\tilde{\mathscr{S}}_{T,T} = ch_T^3$ *for some* $c > 0$.

*Proof.* We use the fact that for any matrix $D : \mathbb{R}^m \to \mathbb{R}^n$ and $u \in \mathbb{R}^n$, we have $\sigma_1 \|u\| \leq \|Du\| \leq \sigma_n \|u\|$ where $\sigma_1, \sigma_n$ are the minimum and maximum singular values of $D$, respectively. Applying this to the Jacobian, we see that for any $\hat{z} \in \hat{T}$

$$\sigma_{\min}\|\hat{x} - \hat{y}\| \leq c_1 h_T \|\hat{x} - \hat{y}\| \leq \|dF_T(\hat{z})(\hat{x} - \hat{y})\| \leq c_2 h_T \|\hat{x} - \hat{y}\| \leq \sigma_{\max}\|\hat{x} - \hat{y}\|. \tag{40}$$

Taking $\hat{z} = t(\hat{x} - \hat{y}) + \hat{y}$ and integrating the previous inequalities for $t \in [0,1]$, by the Fundamental Theorem of Calculus we have

$$c_1 h_T \|\hat{x} - \hat{y}\| \leq \|F(\hat{x}) - F(\hat{y})\| \leq c_2 h_T \|\hat{x} - \hat{y}\|. \tag{41}$$

Therefore,

$$\frac{1}{\|F(\hat{x}) - F(\hat{y})\|} = c \frac{1}{h_T \|\hat{x} - \hat{y}\|}. \tag{42}$$

With all of this, we have for the local operator $\mathscr{S}_{T,T}$

$$\begin{aligned}
\mathscr{S}_{T,T} &= \int_T \int_T \frac{1}{4\pi|x-y|} dS_x dS_y, \\
&= \int_{\hat{T}} \int_{\hat{T}} \frac{1}{4\pi|F(\hat{x}) - F(\hat{y})|} dF_T(\hat{x}) dF_T(\hat{y}) dS_{\hat{x}} dS_{\hat{y}}, \\
&= c \int_{\hat{T}} \int_{\hat{T}} \frac{1}{4\pi|\hat{x}-\hat{y}|} \frac{h_T^2 h_T^2}{h_T} dS_{\hat{x}} dS_{\hat{y}}, \\
&= c h_T^3 |\hat{T}|^2, \\
&= c h_T^3.
\end{aligned} \tag{43}$$

□

If the mesh $\tau$ is *shape regular* (i.e. $h_T = c|T|^{1/2}$ [31]), then the scaling in Equation 20 is clear.

#### S1.4 Proof of Vanishing Surrogate Discrepancy

To make the notion of discrepancy more rigorous, denote by $\tau_l$ the discrete mesh at adaptive iteration $l$ (with $\tau_0$ denoting the initial mesh). We use implicit notation $\rho_l := \rho_{\tau_l}$ to denote the Galerkin solution at adaptive iteration $l$, with similar notation for any other iteration dependent quantities (e.g. charge residual $e_l$). We assume that the meshes are always refined in the adaptive process, that is $\tau_l \subset \tau_{l-1} \subset \cdots \subset S$. Additionally, we assume that the Galerkin solutions are chained into the Sobolev space of the previous mesh, i.e.

$$\rho_l \in \mathscr{H}^{-1/2}(\tau_l) \subset \mathscr{H}^{-1/2}(\tau_{l-1}) \subset \cdots \subset \mathscr{H}^{-1/2}(S). \tag{44}$$

We now define the projection $\mathscr{P}_l : \mathscr{H}^{-1/2}(\tau_l) \to \mathscr{H}^{-1/2}(\tau_{l-1})$ as the operator such that for any $u \in \mathscr{H}^{-1/2}(\tau_l)$,

$$\langle u - \mathscr{P}_l u, v \rangle = 0 \quad \forall v \in \mathscr{H}^{-1/2}(\tau_{l-1}). \tag{45}$$

(Note that by the chaining property, this inner product is well-defined). Now we will define the surrogate function

$$\xi_l := \rho_l - \mathscr{P}_l \rho_l. \tag{46}$$

Notice that $\xi_l$ intrinsically acts as a surrogate for the charge residual since it is the charge solution at iteration $l$ minus the contribution of charge from the previous iteration, as shown below.

**Lemma S1.5.** *The surrogate can also be written as* $\xi_l = \rho_l - \rho_{l-1}$.

*Proof.* $\rho_l$ is the Galerkin solution which solves

$$\langle \rho - \rho_l, v \rangle = 0 \quad \forall v \in \mathscr{H}^{-1/2}(\tau_l) \subset \mathscr{H}^{-1/2}(\tau_{l-1}). \tag{47}$$

By definition, $\mathscr{P}_l \rho_l$ solves

$$\langle \rho_l - \mathscr{P}_l \rho_l, v \rangle = 0 \quad \forall v \in \mathscr{H}^{-1/2}(\tau_{l-1}). \tag{48}$$

Subtracting Equations 47 and 48 yields

$$\langle \rho - \mathscr{P}_l \rho_l, v \rangle = 0 \quad \forall v \in \mathscr{H}^{-1/2}(\tau_{l-1}). \tag{49}$$

But $\rho_{l-1}$ is the Galerkin solution which solves

$$\langle \rho - \rho_{l-1}, v \rangle = 0 \quad \forall v \in \mathscr{H}^{-1/2}(\tau_{l-1}). \tag{50}$$

Hence, $\mathscr{P}_l \rho_l = \rho_{l-1}$. □

We can more easily analyze the discrepancy with this form of the surrogate $\xi_l$.

**Proposition S1.6.** *For the surrogate* $\xi_l$*, we have* $\lim_{l\to\infty} D(\xi_l) = 0$.

*Proof.* First, notice that the charge residual can be written as

$$\begin{aligned} e_l &= \rho - \rho_l, \\ &= (\rho - \rho_{l-1}) - (\rho_l - \rho_{l-1}), \\ &= e_{l-1} - \xi_l. \end{aligned} \tag{51}$$

Particularly, $\xi_l = e_{l-1} - e_l$, and $|||e_l||| \le |||e_{l-1}|||$. Therefore, we have

$$\begin{aligned} |||e_l - \xi_l||| &= |||2e_l - e_{l-1}|||, \\ &\le 2|||e_l||| + |||e_{l-1}|||, \\ &\le 3|||e_{l-1}|||, \\ &\to 0. \end{aligned} \tag{52}$$

□

In practice, we find that $\xi_l$ does not serve well as a direct substitute in the refinement criterion $\eta_T$ (e.g. the electrode currents tend to converge "too early" to incorrect values). This is likely due to the fact that the surrogate function prioritizes triangles with the greatest change in the charge from each iteration, which will invariably occur on and near the electrodes, and hence rarely or never refines the triangles in the deeper tissues. We can reduce the number of refinements which occur on previously refined triangles by prioritizing triangles with a larger area. Hence, we introduce the area scaled surrogate

$$\xi^*_{l,T} = A_{l,T}\xi_l, \tag{53}$$

where $A_{l,T} = |T_l|$, that is, the area of triangle $T$ on iteration $l$. We will use the shorthand notation $\xi^*_l = A_l\xi_l$, as the implicit dependence on the triangles $T$ is clear.

**Proposition S1.7.** *For the surrogate* $\xi^*_l$*, we have* $\lim_{l\to\infty} D(\xi^*_l) = 0$.

*Proof.* The proof follows in the same way as Proposition S1.6, where we now have

$$\begin{aligned} e_l - \xi^*_l &= e_l - A_l\xi_l, \\ &= e_l - A_l(e_{l-1} - e_l), \\ &= (1 + A_l)e_l - A_le_{l-1}. \end{aligned} \tag{54}$$

Since the mesh is always refined, we have the guaranteed existence of $A_{\max} = \max_{l,T\in\tau_l} |T_l|$, the maximum triangle area achieved across all iterations. Then

$$\begin{aligned} |||e_l - \xi^*_l||| &\le |||(1 + A_l)e_l||| + |||A_le_{l-1}|||, \\ &\le (1 + A_{\max})|||e_l||| + A_{\max}|||e_{l-1}|||, \\ &\le (1 + 2A_{\max})|||e_l|||, \\ &\to 0. \end{aligned} \tag{55}$$

□

Note that the proof of Proposition S1.7 is valid under any scaling $A_l^p \xi_l$ with $p \geq 0$. Hence, we can choose various scaling powers of triangle area depending on how much priority we would like to place on refining larger (or smaller) triangles.

Notice that if we were to choose as a surrogate the charge solution at each iteration, then the discrepancy is $D(\rho_l) \to |||\rho|||$, which is not sufficient by our definition. If scaling by the area, then

$$D(A_l \rho_l) \to \lim_{l \to \infty} (\max_{T \in \tau_l}(A_l)) \cdot |||\rho|||. \tag{56}$$

Hence, using $A_l \rho_l$ (or $A_l^p \rho_l$ for $p > 0$) as a surrogate will only have vanishing discrepancy if the maximum triangle area on each iteration tends to zero as well.

### S2 Supplement: Review of BEM-FMM

#### S2.1 Boundary Element Fast Multipole Method

Consider $\Omega$ comprising a volume of interest and $S = S_1 \cup \cdots \cup S_n$ the union of disjoint surfaces (the cortical tissues) within $\Omega$. The charge based formulation of BEM is the integral equation for determining the surface charge density $\rho(\mathbf{x})$ [8, 17, 18]

$$\frac{\rho(\mathbf{x})}{2\varepsilon_0} - \kappa(\mathbf{x})\mathbf{n}(\mathbf{x}) \cdot \int_S \frac{\rho(\mathbf{y})}{4\pi\varepsilon_0} \frac{\mathbf{x}-\mathbf{y}}{|\mathbf{x}-\mathbf{y}|^3} dS(\mathbf{y}) = \kappa(\mathbf{x})\mathbf{E}^i(\mathbf{x}) \cdot \mathbf{n}(\mathbf{x}) \tag{57}$$

where

- $\mathbf{n}(\mathbf{x})$ is the outward unit normal vector at $\mathbf{x} \in S$;
- $\varepsilon_0$ is dielectric permittivity of free space;
- $\kappa(\mathbf{x}) = (\sigma_- - \sigma_+)/(\sigma_- + \sigma_+)$ is the conductivity contrast with respect to inner and outer conductivities $\sigma_-$ and $\sigma_+$, respectively;
- $\mathbf{E}^i(\mathbf{x})$ is the impressed electric field induced by the TES or EEG electrodes.

We assume here that the electric scalar potential $u(\mathbf{x})$ is defined as a single-layer potential of the surface charge density

$$u(\mathbf{x}) = \int_S \rho(\mathbf{y})\Phi(\mathbf{x},\mathbf{y})dS(\mathbf{y}) \tag{58}$$

where $\Phi(\mathbf{x},\mathbf{y}) = 1/(4\pi|\mathbf{x}-\mathbf{y}|)$ is the fundamental solution to the Laplace equation. In this manner, $u(\mathbf{x})$ is a solution to the Dirichlet problem

$$\begin{aligned} \Delta u &= \nabla \cdot \mathbf{J}^i \quad && \Omega \setminus S, \\ u &= f && S. \end{aligned} \tag{59}$$

where $\mathbf{J}^i$ is the impressed current (with $\nabla \cdot \mathbf{J}^i = 0$ outside the outer-most tissue) and $f$ is the boundary potential. Note that $f$ is an a priori unknown quantity; the boundary relation can be adjusted to a condition on the jump in potential across the surfaces which leads to the classical derivation of the potential based BEM (see [18]).

Denote by $\mathscr{S}$ the single-layer potential operator such that $u(\mathbf{x}) = \mathscr{S}(\rho(\mathbf{x}))$, and its corresponding discretized form $\hat{\mathscr{S}}$. By extrapolating to the wider single-layer potential operator $\mathscr{S}$, we can consider $\rho$ as solutions to the general operator equation

$$\hat{\mathscr{S}}\rho = f, \tag{60}$$

whose solution we may approximate via the Galerkin method

$$\langle \hat{\mathscr{S}}\rho_h, v\rangle = \langle f, v\rangle, \tag{61}$$

which is satisfied for any test function $v$ (see Section S1.1). This can be written as a linear system of equations by assuming a finite basis $s_i(\mathbf{x})$ of the surface charge so that

$$\rho_h(\mathbf{x}) = \sum_i \rho_i s_i(\mathbf{x}) \tag{62}$$

for unknown coefficients $\rho_i$. If $S$ is discretized into a triangular mesh, then this summation is taken over all $N$ triangles and the discretized Galerkin approximation of the charge at triangle $T$ can be explicitly calculated as [17, 18]

$$\rho_T = 2\varepsilon_0\kappa(\mathbf{x}_T)\mathbf{n}(\mathbf{x}_T)\cdot\left(\sum_{K\neq T}\frac{|K|\rho(\mathbf{x}_K)(\mathbf{x}_T-\mathbf{x}_K)}{4\pi\varepsilon_0|\mathbf{x}_T-\mathbf{x}_K|}+\mathbf{E}^i(\mathbf{x}_T)\right) \tag{63}$$

where $|K|$ denotes the area of triangle $K$, $\mathbf{x}_T$ and $\mathbf{x}_K$ the center points of triangles $T$ and $K$, respectively, and the calculation of the impressed electric field is accelerated with FMM.

The Galerkin residual $r = f - \hat{\mathscr{S}}\rho_h$ is used as an estimator for the convergence of the iterative Galerkin method. In classic adaptive BEM theory, this residual serves as the basis for mesh refinement criteria [12]. Since the boundary data $f$ is unknown, directly applying the classical theory of adaptive BEM is challenging. We can approximate the Galerkin residual by $f' - \hat{\mathscr{S}}\rho_h$, where $f'$ is discrete boundary data determined by constructing a preconditioner matrix on the electrode-scalp interface (see Section 2.2 of the main paper). However, we found that in practice, using this approximation of the residual resulted in inferior adaptive mesh refinement results when compared to other refinement criteria. Hence our desire to derive a different criterion, see Supplement S1.

### S3 Supplement: Additional Tables and Figures

Table 1: Table of tissue conductivities for the three model types analyzed. The third column shows the (inner) tissue conductivity, and the fourth column shows the designated outer tissue. Note that for all tissues in the Sim4Life model, we designate the outer tissue as Freespace for purely notational purposes; the tissues are non-manifold, and thus no outer tissue can be specified. The tissue conductivities used for the Sim4Life models can be found in [32].

| Model | Tissue | Conductivity (Inner) | Outer Tissue |
|---|---|---|---|
| Sphere | $S_1$ | 0.465 | Freespace |
| | $S_2$ | 0.01 | $S_1$ |
| | $S_3$ | 1.654 | $S_2$ |
| | $S_4$ | 0.275 | $S_3$ |
| | $S_5$ | 0.126 | $S_4$ |
| SimNIBS | Skin | 0.465 | Freespace |
| | Bone | 0.01 | Skin |
| | CSF | 1.654 | Bone |
| | GM | 0.275 | CSF |
| | WM | 0.126 | GM |
| | Eyes | 1.2 | Skin |
| | Ventricles | 1.654 | WM |
| Sim4Life | Skin | 0.148 | Freespace |
| | Air_internal | 0 | Freespace |
| | Amygdala | 0.419 | Freespace |
| | Artery | 0.662 | Freespace |
| | Brainstem | 0.348 | Freespace |
| | Cartilage | 0.739 | Freespace |
| | Caudate_nucleus | 0.348 | Freespace |
| | Cerebellum_grey_matter | 0.419 | Freespace |
| | Cerebellum_white_matter | 0.348 | Freespace |
| | Cerebrospinal_fluid | 1.88 | Freespace |
| | Cerebrum_grey_matter | 0.419 | Freespace |
| | Cerebrum_white_matter | 0.348 | Freespace |
| | Dura | 0.06 | Freespace |
| | Eyes | 2.16 | Freespace |
| | Globus_pallidus | 0.348 | Freespace |
| | Hippocampus | 0.419 | Freespace |
| | Intervertebral_disc | 0.739 | Freespace |
| | Midbrain_ventral | 0.348 | Freespace |
| | Mucosa | 0.461 | Freespace |
| | Muscle | 0.461 | Freespace |
| | Muscle_ocular | 0.461 | Freespace |
| | Nasal_septum | 0.175 | Freespace |
| | Nerve_cranial_II_optic | 0.348 | Freespace |
| | Nucleus_accumbens | 0.419 | Freespace |
| | Other_tissues | 0.461 | Freespace |
| | Parotid_gland | 0.481 | Freespace |
| | Putamen | 0.348 | Freespace |
| | Skull_cancellous | 0.0805 | Freespace |
| | Skull_cortical | 0.0063 | Freespace |
| | Spinal_cord | 0.611 | Freespace |
| | Sublingual_gland | 0.481 | Freespace |
| | Submandibular_gland | 0.481 | Freespace |
| | Tendon_galea_aponeurotica | 0.368 | Freespace |
| | Tendon_temporalis | 0.368 | Freespace |
| | Thalamus | 0.475 | Freespace |
| | Tongue | 0.461 | Freespace |
| | Vein | 0.662 | Freespace |
| | Ventricles | 1.88 | Freespace |
| | Vertebrae_cancellous | 0.0805 | Freespace |
| | Vertebrae_cortical | 0.0063 | Freespace |

Table 2: Table of the Sim4Life tissues used 40-tissue head models. Cell color corresponds to the tissue segment color depicted in Fig. 3b of the main paper.

| | |
|---|---|
| SKIN | BRAINSTEM |
| AIRINTERNAL | CARTILAGE |
| OTHERTISSUES | CAUDATE_NUCLEUS |
| BONEC | CEREBELLUM_GREY_MATTER |
| BONET | CEREBELLUM_WHITE_MATTER |
| DURA | GLOBUS_PALLIDUS |
| CSF | INTERVERTEBRAL_DISC |
| GM | MIDBRAIN_VENTRAL |
| WM | MUSCLE |
| EYES | NASAL_SEPTUM |
| MUCOSA | NUCLEUS_ACCUMBENS |
| MUSCLEOCULAR | PAROTID_GLAND |
| SPINALCORD | PUTAMEN |
| ARTERY | SUBLINGUAL_GLAND |
| VEIN | SUBMANDIBULAR_GLAND |
| NERVE | TENDON_GALEA_APONEUROTICA |
| HIPPOCAMPUS | TENDON_TEMPORALIS |
| THALAMUS | TONGUE |
| VENTRICLES | VERTEBRAE_CANCELLOUS |
| AMYGDALA | VERTEBRAE_CORTICAL |
